# CD248 activates TGF-β receptor I to promote vascular remodeling in pulmonary arterial hypertension

**DOI:** 10.64898/2026.04.22.720270

**Authors:** Luke I Jones, Hannah J McIntire-Ray, Angela N Morales, Shia Vang, Meghan J Hirsch, Andres J Gonzalez Coba, Emma Lea Matthews, Katherine L Adriatico, Nicholas P Harris, Iram Zafar, Dongqi Xing, Vivian Y Lin, Liping Tian, Gregory A Payne, Aftab Ahmad, Raed A Dweik, J Michael Wells, Heather M Olson, Jennifer E Kyle, Geremy C Clair, Stefanie Krick, Jarrod W Barnes

**Affiliations:** Medical Scientist (MD-PhD) Training Program, University of Alabama at Birmingham, Birmingham AL USA; Department of Medicine, Division of Pulmonary, Allergy & Critical Care Medicine, University of Alabama at Birmingham, Birmingham AL USA; Department of Anesthesia and Perioperative Medicine, University of Alabama at Birmingham, Birmingham AL USA; Department of Pulmonary, Allergy and Critical Care Medicine, Respiratory Institute, Cleveland Clinic, Cleveland OH USA; Department of Medicine, Division of Cardiovascular Diseases, University of Alabama at Birmingham, Birmingham AL USA; Lung Health Center, University of Alabama at Birmingham, Birmingham AL USA; Biological Sciences Division, Pacific Northwest National Laboratory, Richland, WA USA

**Keywords:** CD248, TGF-β signaling, TGFβ receptor I, TβRI, vascular remodeling, pulmonary arterial hypertension, smooth muscle cell, cell proliferation, extracellular matrix, Ontuxizumab, mass spectrometry

## Abstract

I.

**Background:** Pulmonary arterial hypertension (PAH) is a debilitating cardiopulmonary disease characterized by progressive remodeling of the pulmonary vasculature. Pathologic transforming growth factor-β (TGF-β) signaling is an essential driver of vascular remodeling in PAH. While global inhibitors of TGF-β exist, their clinical application is limited by systemic adverse effects. Therefore, a critically unmet need in PAH is to identify pulmonary vascular-specific regulators of the TGF-β axis, which would selectively enhance clinical efficacy while minimizing adverse effects. As the clinical care of PAH largely promotes vasodilation, and only one FDA-approved agent targets vascular remodeling, this study aimed to identify selective, therapeutically targetable regulators of the TGF-β axis in the PAH pulmonary vasculature.

**Methods:** CD248 was identified via liquid chromatography-tandem mass spectrometry (LC-MS/MS) proteomics in human lungs. CD248 levels were assessed across human, rat, and mouse lung tissues using western blotting, RTqPCR, and/or immunofluorescence techniques. CD248-null (CD248^-/-^) mice were used to study the contribution of CD248 to hypoxia-sugen (H/S)-induced PAH. The mechanistic role of CD248 in PAH vascular remodeling and TGF-β signaling was assessed by genetic (siRNA knockdown; overexpression) and pharmacologic (Ontuxizumab) manipulation of primary human pulmonary vascular cells.

**Results:** LC-MS/MS proteomics coupled with pathway enrichment analysis of human lung tissue identified CD248 as a putative mediator of vascular remodeling that is elevated in PAH lungs. CD248 was elevated in PAH pulmonary artery smooth muscle cells (PASMCs) across human, rat, and mouse lung tissue. CD248^-/-^ mice were protected from H/S-induced elevations in right ventricular (RV) systolic pressure (RVSP), RV hypertrophy, and pulmonary artery muscularization. CD248 knock-down reduced cell proliferation and migration of primary PAH PASMCs. CD248 was essential for phospho-activation of TGF-β receptor I (TβRI) at S165 and canonical phosphorylation of SMAD3 at S423/425. CD248 loss blunted TGF-β-induced gene expression (FN1, Col1α1, α-SMA) and activated expression of the vasoprotective matrix metalloprotease, MMP-8. Mechanistically, CD248 interacted with and enhanced *de novo* phosphorylation and stability of TβRI, blocking its ubiquitin-mediated proteasomal degradation. Ontuxizumab promoted TβRI instability and attenuated the production of FN1, Col1α1, and α-SMA in primary PAH PASMCs.

**Conclusions:** This work identifies CD248 as a previously unrecognized co-activator of TβRI in PAH. As CD248 is largely quiescent in most adult tissues yet pathologically upregulated in the PAH pulmonary vasculature, this study supports the potential of anti-CD248 therapy as a novel pulmonary vascular-specific alternative to systemic TGF-β inhibition.

## II. INTRODUCTION

Pulmonary arterial hypertension (PAH) is a devastating cardiopulmonary disease with a significant 40% 5-year mortality rate^1^. Within the PAH lung, structural and functional remodeling of the distal pulmonary arteries results in luminal narrowing, stiffening, and eventual obliteration of the precapillary circulation. Elevated pulmonary vascular resistance results in hemodynamic overload in the right heart and eventual right ventricular failure, the leading cause of mortality^1–3^. The clinical care of PAH patients remains supportive and current therapies largely promote vasodilation as only one FDA-approved therapy targets vascular remodeling^4–7^.

PAH vasculopathy is characterized by concentric neointimal and medial hyperplasia, apoptosis resistance, and aberrant deposition of extracellular matrix (ECM) within the pulmonary vascular wall^8^. More specifically, arterial muscularization involves the transition of pulmonary arterial smooth muscle cells (PASMCs) from quiescence to a pro-remodeling phenotype^9^. The bone morphogenetic protein (BMP) subfamily of the TGF-β superfamily has been implicated in this transition as mutations in the anti-proliferative BMP type II receptor (BMPR2) are reported in up to 75% of hereditary PAH cases^9–12^. Abundant evidence now suggests that TGF-β itself may be a primary driver of vascular remodeling in idiopathic PAH^11, 13–16^. In order to inform PAH therapeutic development, it is critical to understand the molecular regulation of TGF-β-induced pulmonary vascular remodeling.

TGF-β signaling is vital for cell growth, differentiation, and inflammation, making this axis indispensable for vascular homeostasis^17–19^. Interestingly, TGF-β receptor I (TβRI) is highly expressed in human PAH pulmonary arteries^20^ and selective inhibition of TβRI has been shown to attenuate PASMCs proliferation^21^. TβRI is also elevated in the monocrotaline (MCT) rat model of PAH^22^, and its inhibition was found to protect against and even reverse MCT-induced disease^21, 23^. These data suggest that TGF-β signaling, specifically TβRI, is a viable therapeutic target in PAH^13^; however, systemic TGF-β superfamily inhibition with sotatercept^4^ has produced notable adverse effects in PAH clinical trials^19^. Therefore, pulmonary vascular-selective molecular regulators of TGF-β signaling have great therapeutic potential in PAH, but it was previously unknown whether such mechanisms exist. Here, we describe that CD248 (Endosialin), a developmental glycoprotein that is quiescent in most adult tissues but pathologically reactivated in the PAH pulmonary vasculature, functions as a novel co-activator and stabilizer of TβRI.

CD248 is a transmembrane, heavily sialylated glycoprotein of the C-type lectin-like (CTL) domain superfamily. CD248 is critical for embryonic body patterning and vascular development; however, its expression is largely repressed in non-pathologic tissues throughout adulthood^24, 25^. Extensive literature has demonstrated that pathologic reactivation of CD248 in fibrotic disease and malignancy enhances cell proliferation, migration, and ECM remodeling^26–36^. TGF-β signaling may even provide a mechanistic link between CD248 and these processes^30^. Interestingly, CD248 has been proposed as a novel candidate of lung vascular remodeling in heritable, BMPR2 mutation-negative PAH patients^37^, yet the contribution of CD248 to PAH pathogenesis remains undefined.

This work identifies CD248 as a previously unrecognized co-activator of TβRI in human PAH PASMCs. We report that genetic loss of CD248 attenuates hypoxia-sugen (H/S)-induced PAH in mice and that the anti-CD248 clinical therapy, Ontuxizumab^38–40^ reduces hallmarks of vascular remodeling in primary human PAH PASMCs. Mechanistically, CD248 interacts with and is essential for the stability and *de novo* phosphorylation of TβRI. This study supports the potential for anti-CD248 therapy as a pulmonary vascular-specific alternative to systemic TGF-β inhibition.

## III. RESULTS

### Proteomic analysis identifies signatures of vascular remodeling in human PAH lungs

To identify novel putative targets in the PAH lung, we performed LC-MS/MS proteomics on 6 PAH and 6 control (CTRL) human subjects (**Fig 1a**, **Supplemental Fig 1**). Principal component analysis revealed clear separation between CTRL and PAH patient lungs with PC1 and PC2 accounting for 26.14% and 17.5% of total variance, respectively, suggesting disease-driven proteomic differences (**Fig 1b**). LC-MS/MS outputs were filtered for statistical significance (p<0.05) and additional analysis was carried forward on outputs greater than 1.5-fold change (FC) between groups. Of the 6,817 proteins detected, 220 differentially expressed proteins (DEPs) were identified in PAH (**Fig 1c**). 77 DEPs were positively regulated (pDEPs) and 143 were negatively regulated (nDEPs) **(Fig 1d)**. A complete list of DEPs can be found in **Supplemental Table 1**. Immunoblotting was used to validate select LC-MS/MS-identified DEPs in patient-matched lung tissue (**Supplemental Fig 2a,b**). A pathway enrichment analysis of all DEPs was then performed via Metascape^41^. The top 18 PAH-enriched pDEP pathways are shown in **Fig 1e** (a comprehensive list is provided in **Supplemental Fig 3**). Identified pathways include DEPs linked to vascular remodeling processes such as extracellular matrix (ECM) remodeling, cell growth, and TGF-β responsiveness, many of which are well-established markers of pathologic vascular remodeling in PAH (yellow diamonds; **Fig 1d; Supplemental Figure 2c**).

**Figure 1.**
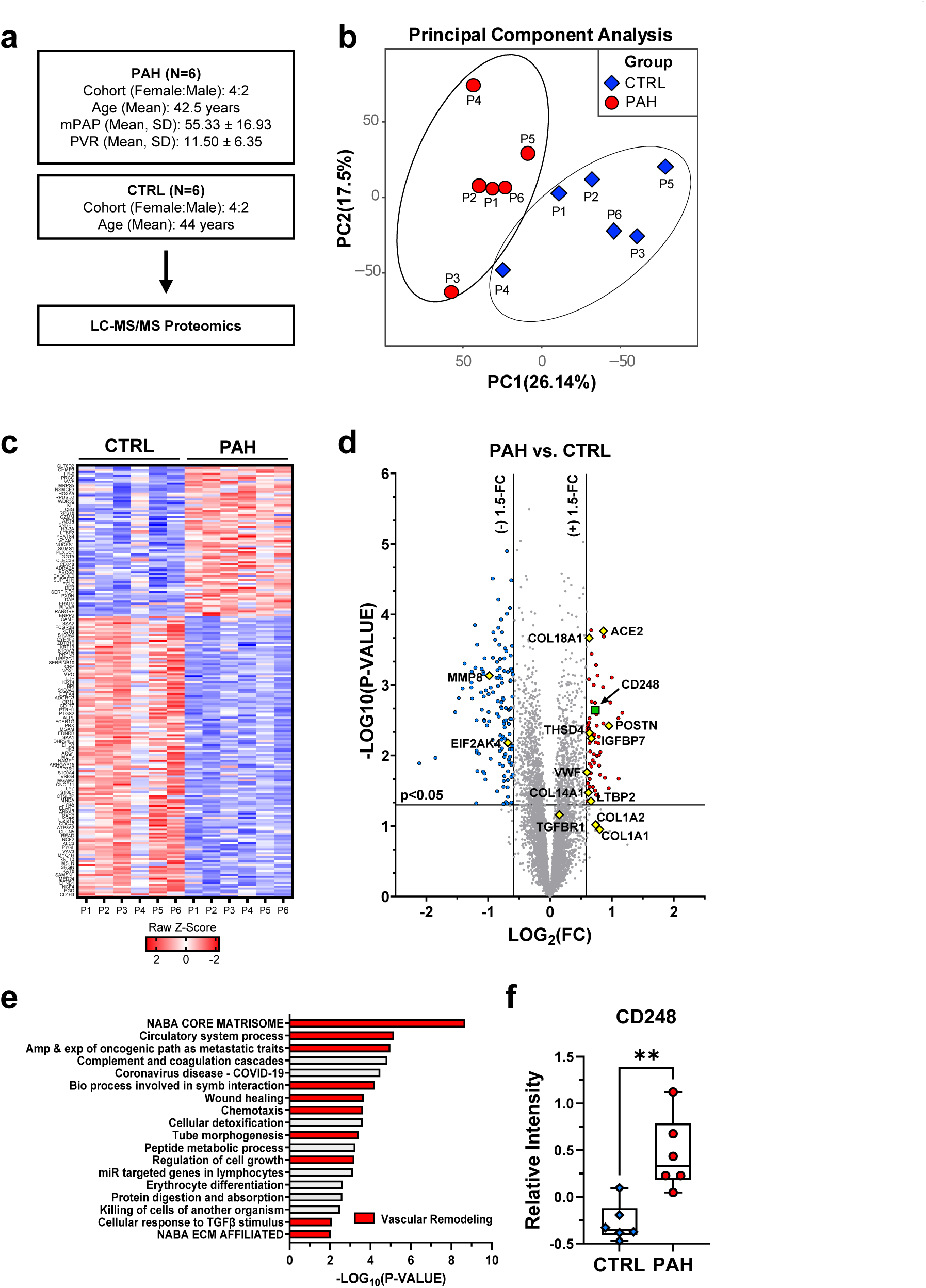
CD248 is elevated in the PAH lung. Liquid chromatography 145 tandem mass spectrometry (LCMS/MS) was performed on lung tissue collected from 12 human subjects (6 PAH, 6 CTRL). **(A)** Overview of human cohort demographics and clinical metrics at lung collection. Mean pulmonary artery pressure (mPAP, mmHg) and pulmonary vascular resistance (PVR, Wood units) are reported as mean ± SD; N=6 human subjects. **(B)** Principal component analysis of CTRL (red circle) and PAH (blue diamond) human lungs. **(C)** Heat map of Z-scores for differentially expressed proteins (DEPs) in PAH vs. CTRL lungs. Positively regulated proteins (pDEPs) in PAH are indicated in red; negatively regulated proteins are indicated in blue (nDEPs). Individual patients are labeled P1-P6 for each group and represented by columns. **(D)** Volcano distribution of proteins from **B**; significance was determined as P<0.05 and ≥ ±1.5-fold change (FC). Vascular remodeling (VR)-associated proteins are indicated by yellow diamonds; CD248 is indicated by a green square. **(E)** Metascape41 pathway enrichment analysis of all pDEPs from PAH lung tissue; the top 18 pathways are shown. VR-associated processes are indicated in red. **(F)** Mean-centered relative intensity of CD248 in PAH lungs relative to CTRL. N=6 human subjects per group. *P<0.05, 158 **P<0.01, ***P<0.001. Statistical analyses were performed using ANOVA and two-tailed heteroscedastic Student’s t-test to compare the experimental groups (CTRL and PAH).

### CD248 is elevated in pulmonary artery smooth muscle cells of human lungs and pre-clinical murine models of PAH

CD248 was identified to be elevated in the PAH lung via MS proteomics (**Fig 1f**). To validate this expression pattern, we performed a Western blot (WB) analysis of patient-matched lung tissue; these data confirmed elevated CD248 levels in PAH relative to controls (**Fig 2a**). CD248 was found to localize to the PAH pulmonary vasculature, notably within α-smooth muscle actin-positive (α-SMA+) pulmonary artery smooth muscle cells (PASMCs) via IF (**Fig 2b**). We further evaluated CD248 expression in an additional cohort of primary human pulmonary vascular cells, demonstrating a PASMC-specific increase (**Fig 2c**) with no corresponding change in PAECs (**Fig 2e**). Reanalysis of a separate PAH PASMCs cohort^42^ also identified >4-fold higher CD248 mRNA levels in PAH PASMCs versus controls (**Fig 2d**; GSE144274). Interestingly, circulating CD248 levels were significantly reduced in PAH human plasma (**Fig 2f**), supporting a PASMC-specific upregulation of CD248 expression in the PAH lung that is independent from systemic circulation. In the non-pathologic adult lung, single-cell RNA sequencing data suggest that CD248 expression is largely quiescent, yet limited expression can be identified in vascular smooth muscle cells and fibroblasts^43^. Complementing these human data, we characterized the expression of CD248 in the lung of two well-described PAH murine models: the hypoxia-SU5416 (H/S) rat and mouse^44^. CD248 was significantly elevated in the H/S rat lung (**Fig 2g**), notably in α-SMA+ PASMCs (**Fig 2h**). In addition, a near 2-fold increase in CD248 levels was observed in the H/S mouse lung (**Fig 2i**), where its abundance was also observed in α-SMA+ PASMCs (**Fig 2j**). These data collectively suggest that elevated CD248 expression within α-SMA+ PASMCs is a conserved phenomenon across both human subjects and established rodent models of PAH.

**Figure 2.**
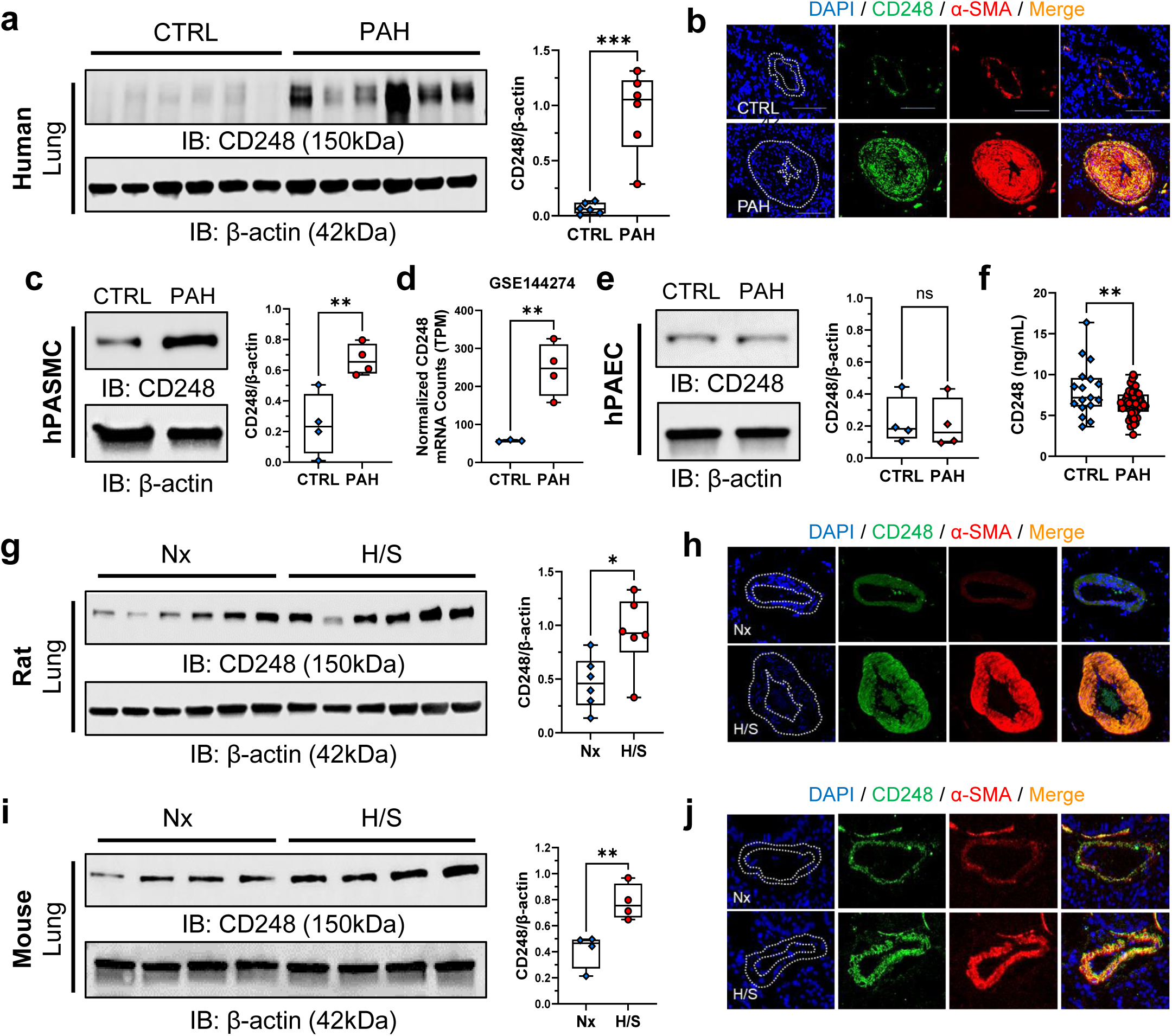
CD248 is elevated in pulmonary artery smooth muscle cells of human and murine models of PAH. **(A)** Western blot (WB) for CD248 in CTRL and PAH human lung tissue matched to **Fig 1**; densitometric calculations are reported (N=6 per group). **(B)** Immunofluorescence (IF) imaging of CD248 (green) and α-smooth muscle actin (α-SMA; red) in CTRL and PAH lung tissue. Merged staining is indicated in orange, and DAPI (blue) was used as a nuclear stain. Dotted lines outline the vascular wall. **(C)** Representative WB for CD248 in primary human pulmonary artery smooth muscle cells (PASMCs) with densitometric quantifications; N=4 human subjects per group. **(D)** Normalized CD248 mRNA counts (transcripts per million; TPM) from control and PAH PASMCs (N=3-4 human subjects per group); data obtained from **GSE144274**. **(E)** Representative WB for CD248 in primary human pulmonary artery endothelial cells (PAECs) with densitometric quantifications; N=4 human subjects per group. **(F)** Soluble CD248 levels in PAH plasma relative to controls, determined via ELISA. N=20 CTRL; N=61 PAH. **(G)** WB for CD248 in normoxia (Nx; room air) and 3-week hypoxia-sugen (H/S) rats (see *Methods*) with densitometric quantifications; N=6 animals per group. **(H)** IF imaging of CD248 (green) and α-SMA (red) in Nx and H/S rat lung tissue. Merged staining is indicated in orange, and DAPI (blue) was used as a nuclear stain. Dotted lines indicate the vascular wall. **(I)** WB for CD248 in Nx and 3-week hypoxia-sugen (H/S) mice (see *Methods*) with densitometric quantifications; N=4 animals per group. **(J)** IF imaging of CD248 (green) and α-SMA (red) in Nx and H/S mouse lung tissue. Merged staining is indicated in orange, and DAPI (blue) was used as a nuclear stain. Dotted lines indicate the vascular wall. All WBs are reported as a fractional ratio relative to β-actin as a loading control. *P<0.05, **P<0.01; ns=not significant. Statistical analyses were performed using independent t-tests. A Mann-Whitney test was performed for nonparametric data.

### CD248 knock-out attenuates H/S-induced PAH in mice

We utilized wild-type (WT) and CD248 germline knock-out (KO; −/−) mice to determine the impact of CD248 loss on the development of H/S-induced PAH *in vivo* (**Fig 3a**). After confirming CD248 loss at genetic (**Fig 3b**) and protein (**Fig 3c**) levels, we assessed right ventricular systolic pressures (RVSP) and RV hypertrophy (RV/LV+sep) of all groups following 3-week H/S or normoxia (Nx; room air) exposure. CD248^−/−^ mice exposed to H/S had significantly lower RVSP relative to age– and sex-matched WT mice of the same condition (**Fig 3d**). Similarly, CD248 loss protected against H/S-induced RV hypertrophy (**Fig 3e**). Neither RVSP nor RV hypertrophy were affected by loss of CD248 in Nx control mice. These findings indicate that CD248 is necessary for the development of H/S– induced PAH but is distinct from general cardiopulmonary function in control mice. In addition, H/S-exposed CD248^−/−^ mice had significantly reduced pulmonary artery concentric muscularization relative to WT H/S-exposed mice (**Fig 3f-g**). These data suggest that CD248 loss protects against H/S-induced PAH in mice and is therefore vital to murine PAH pathogenesis.

**Figure 3.**
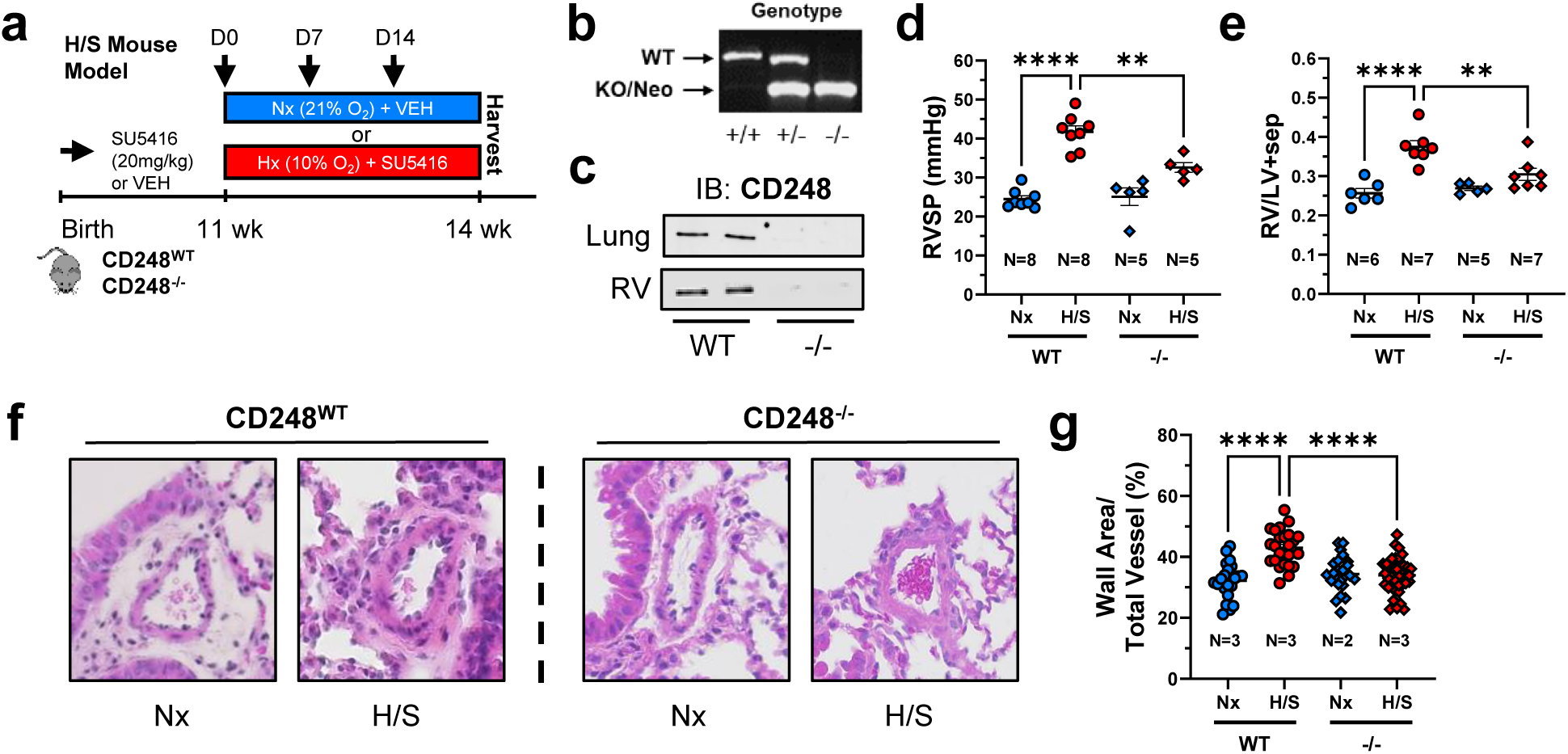
CD248 knock-out attenuates H/S-induced PAH in mice. **(A)** Schematic depicting the timeline for generation of H/S mice. 10-12-week C57BL/6 wild-type (WT) or CD248 germline knock-out (KO; CD248−/−) mice were subjected to either normoxia (Nx; 21% O_2_) or hypoxia (10% O_2_) and delivered a subcutaneous injection of DMSO (vehicle) or SU5416 (20mg/kg) once weekly. All assessments were performed at endpoint (3 weeks). **(B)** PCR identification of CD248 genotype: WT (+/+), Heterozygous (+/-), and knock out (−/−). **(C)** WB for CD248 in WT and −/− mouse lung and right ventricle (RV). **(D)** 3-week right ventricular systolic pressure (RVSP) of anesthetized mice according to the indicated groups; N=5-8 animals per group. **(E)** Fulton index (FI; ratio of RV to LV+septum weight) for all indicated mouse groups (N=5-7 mice per group). **(F)** Representative H&E images of pulmonary arteries (A) from WT and CD248−/− mice exposed to Nx or H/S (average vessel size <150μm). **(G)** Quantification of the vascular wall area relative to total wall area. A mean of 10 vascular fields/per mouse were analyzed. Data are reported as mean ± SEM; *P<0.05, **P<0.01, ****P<0.0001. Analyses were performed via two-way ANOVA with Tukey-Kramer post-hoc comparisons tests.

### CD248 loss attenuates the proliferation and migration of PAH PASMCs

As loss of CD248 attenuated arterial muscularization *in vivo*, we next investigated the impact of CD248-deficiency on the proliferation of primary isolated PASMCs from PAH donors *in vitro*. PAH PASMCs exhibited a 4-fold increase in proliferating cell nuclear antigen (PCNA) expression relative to control PASMCs at baseline (**Fig 4a**), consistent with prior studies^45–47^. 48-hours post-CD248 siRNA knockdown (KD) across 4 patient-derived PAH PASMCs donors, we observed a significant reduction in PCNA levels (**Fig 4b-d**). In addition, genomic incorporation of the Click-iT® AlexaFluor-488 conjugated nucleotide analog, EdU, a surrogate of DNA synthesis and cell proliferation, was also decreased following CD248 KD in PAH PASMCs (**Fig 4e**). These findings were corroborated by a reduction in proliferative metabolic potential (MTT absorbance; **Fig 4f**) and blunted migration/wound repair (**Fig 4g**) following CD248 loss. Collectively, these data indicate that CD248 is vital for PASMC proliferation and migration in PAH.

**Figure 4.**
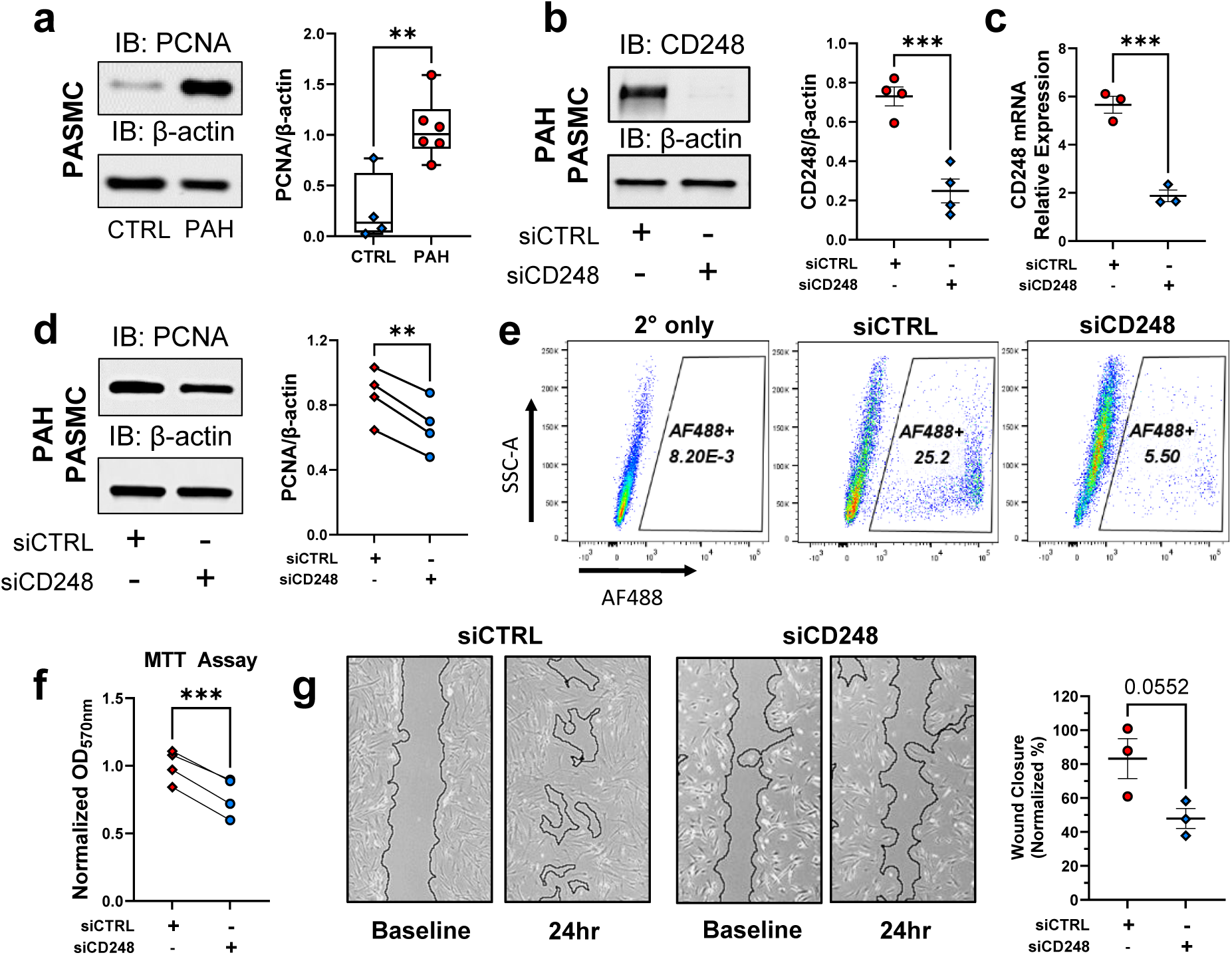
CD248 knock-down reduces the proliferation and migration of PAH PASMCs. **(A)** Representative WB for proliferating cell nuclear antigen (PCNA) in CTRL and PAH PASMCs with densitometric quantification (N=4-6 human donors). **(B)** Representative WB confirmation of CD248 protein knock-down (KD) in PAH PASMCs with CD248-targeted and –nontargeted siRNA (N=4 biological replicates). Cells were transfected overnight with siCTRL or siCD248 then allowed 48 hours of outgrowth before collection. **(C)** Relative CD248 mRNA expression following CD248 siRNA KD in PAH PASMCs (N=3 biological replicates). **(D)** Representative WB for PCNA in PAH PASMCs with and without CD248-targeted siRNA (N=4 human donors). **(E)** Alexa Fluor (AF)-488-EdU incorporation (nucleotide analog) in PAH PASMCs. Cells were transfected as in **B** then stimulated to proliferate with 20% FBS. Gating for AF488+ (proliferative) cells (**Supplemental Figure 4**) is reported as a bolded percentage. A secondary-only negative control is indicated. **(F)** Normalized MTT viability assay in PAH PASMCs transfected with and without CD248-targeted siRNA (N=4 human donors). Following KD, cells were incubated for 2 hours with MTT reagent followed by solubilization with DMSO; absorbance was measured via OD_570_. **(G)** Wound healing scratch assay in CD248 KD PAH PASMCs. Following CD248 KD, a scratch wound was generated across a confluent cell layer. Images were captured immediately following scratch (baseline) and 24 hours post-wound closure. Wound closure was reported as a normalized percentage relative to siCTRL (N=3 biological replicates). Data are reported as mean ± SEM; **P<0.01, ***P<0.001. All WBs are reported as a fractional ratio relative to β-actin as a loading control. Statistical analyses were performed via independent **(A-C, G)** and paired **(D,F)** t-tests. A Mann-Whitney test was performed for nonparametric data.

### CD248 is crucial for phospho-activation of TβRI at S165

We next determined the role of CD248 in canonical TGF-β signaling. Following recombinant TGF-β1 stimulation, we assessed TβRI abundance and phosphorylation as well as downstream signaling transduction and transcriptional regulation of known TGF– β-responsive genes. Subsequent to CD248 KD and acute TGF-β stimulation, we discovered a marked reduction in the phospho-activation of TβRI (**Fig 5a**). This difference was driven by a striking reduction in phosphorylation of TβRI at S165 (**Fig 5b**); however, we also observed a consistent baseline decrease in total TβRI levels following CD248 loss (**Fig 5c**). In addition, phosphorylation of downstream TβRI-target and transcription factor, mothers against decapentaplegic homolog 3 (SMAD3) at S423/425, was significantly reduced following CD248 loss (**Fig 5d**). These findings indicate CD248 deficiency attenuates aberrant TβRI activation and signaling transduction, particularly by reducing TβRI S165 phosphorylation in primary human PAH PASMCs.

**Figure 5.**
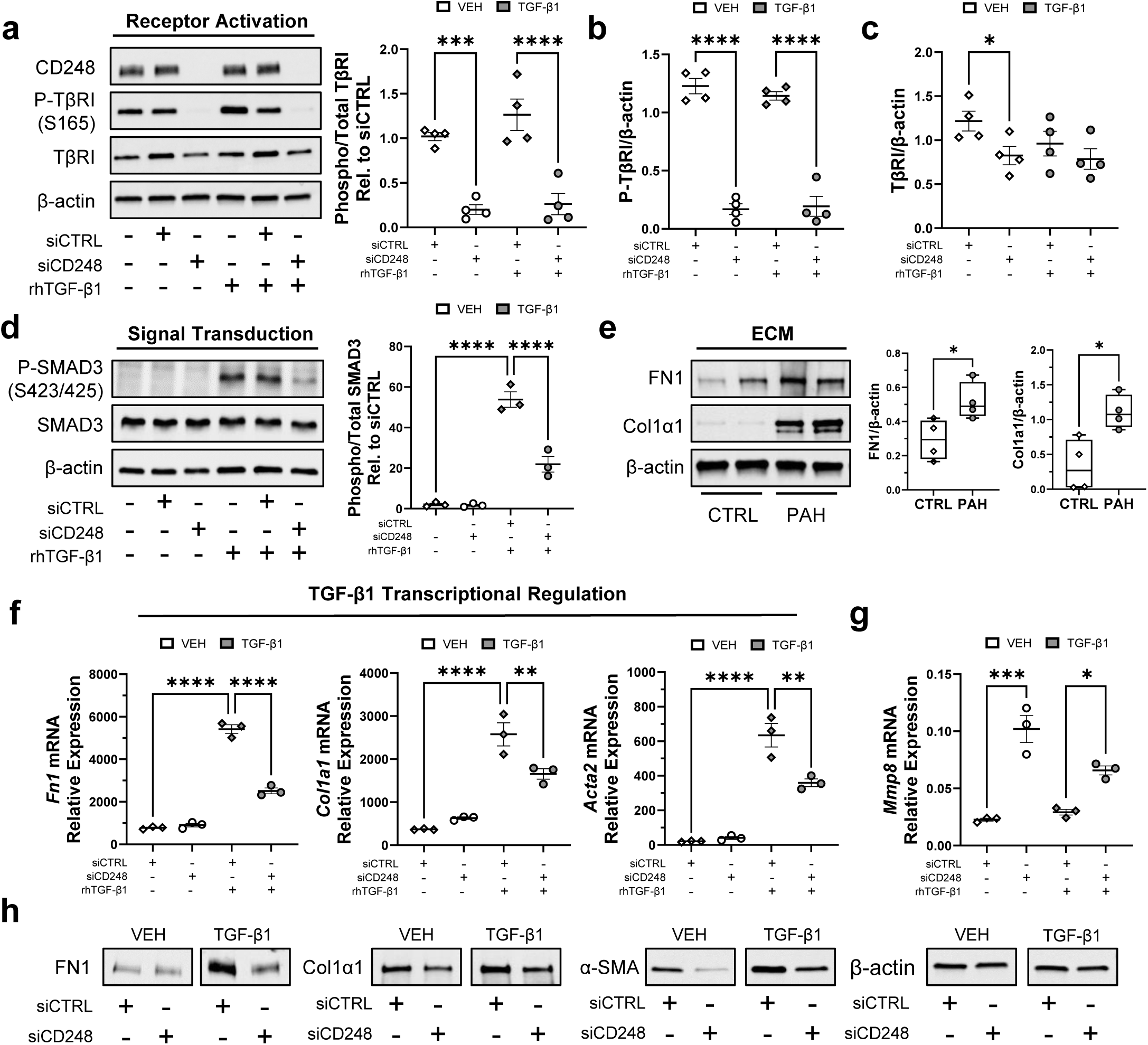
CD248 is critical for TβRI activation and TGF-β-induced vascular remodeling. **(A)** Representative WBs for CD248, P-TβRI (S165), and TβRI in CD248 KD PAH PASMCs. Cells were transfected according to **Fig 4B** then grown for 48 hours followed by serum starvation and acute stimulation with rhTGF-β1 (10ng/mL) for 30 minutes. The densitometric ratio of phosphorylated to total TβRI is reported (N=4 biological replicates). **(B)** Densitometric analysis of P-TβRI from **A. (C)** Densitometric analysis of TβRI from **A**. **(D)** Representative WB for P-SMAD3 (S423/425) and SMAD3 in CD248 KD PAH PASMCs. Cells were transfected and ligand-stimulated as in **A** prior to collection (**Supplemental Figure 5**); N=3 biological replicates. **(E)** Representative WB for vascular extracellular matrix (ECM) proteins fibronectin 1 (FN1) and collagen 1α1 (Col1α1) in control and PAH PASMCs (N=4 human donors per group). All cells were harvested at 80-90% confluence and plates were scraped to include ECM products prior to protein isolation. **(F)** Relative mRNA expression of TGF-β responsive genes *Fn1*, *Col1a1*, and *Acta2* (α-SMA) following CD248 KD and TGF-β stimulation in PAH PAMSCs (N=3 biological replicates). Following CD248 KD, cells were stimulated with rhTGFβ1 (10ng/mL) for 48 hours then harvested. **(G)** Relative mRNA expression of matrix metalloprotease 8 (*Mmp8*) following CD248 KD and TGF-β stimulation as in **F** (N=3 biological replicates). **(H)** Representative WB for vascular ECM proteins, FN1, Col1α1, and α-SMA in PAH PASMCs. As in **F**, cells were transfected with siRNA followed by 48-hour stimulation with rhTGF-β1. Data are reported as mean ± SEM; *P<0.05, **P<0.01, ***P<0.001, ****P<0.0001. All WBs are reported 309 as a fractional ratio relative to β-actin as a loading control. Statistical analyses were performed via independent t-test **(E)** and ordinary 2-way ANOVA with Fisher’s LSD post-hoc comparisons tests **(A-D, F-G).**

### CD248 loss promotes extracellular matrix remodeling

As TGF-β signaling and ECM deposition/remodeling proteins FN1 and Col1α1 are notably elevated in the PAH lung (**Supplemental Fig 6a**) and our primary human PASMC cohort (**Fig 5e**), we evaluated whether CD248 loss was sufficient to reduce the expression of TGF-β-responsive genes. Following CD248 KD in PAH PASMCs, we observed a decline in the transcription of TGF-β1-responsive genes, fibronectin 1 (*Fn1*), collagen 1α1 (*Col1a1*), and alpha smooth muscle actin (*Acta2*) (**Fig 5f**). Interestingly, the vasoprotective matrix metalloprotease-8 (*Mmp8*), which was reduced at baseline in the PAH lung (**Fig 1e**), was transcriptionally activated by CD248 loss (**Fig 5g**). These mRNA transcript findings were confirmed at the protein level as CD248 KD attenuated baseline Col1α1 and α-SMA levels as well as TGF-β-induced FN1, Col1α1, and α-SMA levels in PAH PASMCs (**Fig 5h**). Altogether, these data support an essential role for CD248 as a modulator of pulmonary vascular ECM deposition and remodeling, potentially via activation of TβRI.

### CD248 interacts with and stabilizes TβRI

As CD248 and TβRI are both transmembrane proteins, we sought to determine whether CD248 interacts with TβRI in PAH PASMCs. We performed an endogenous immunoprecipitation (IP) of both TβRI and CD248, which resulted in reciprocal pulldown of both proteins (**Fig 6a-b**). To confirm the specificity of these co-IP results, we repeated the same experiment using epitope-tagged proteins (CD248-FLAG; TβRI-HA) in HEK293T cells, validating the interaction between CD248 and TβRI (**Fig 6c**). We then evaluated the impact of CD248 on TβRI protein stability by implementing a cycloheximide (CHX)-chase experiment to determine the rate of TβRI degradation in CD248-deficient cells. Following CD248 KD (**Fig 6d,e**), the rate of TβRI degradation was markedly enhanced relative to controls (**Fig 6f**). These data suggest that the physical interaction between CD248 and TβRI may be essential for TβRI stability and abundance in PAH.

**Figure 6.**
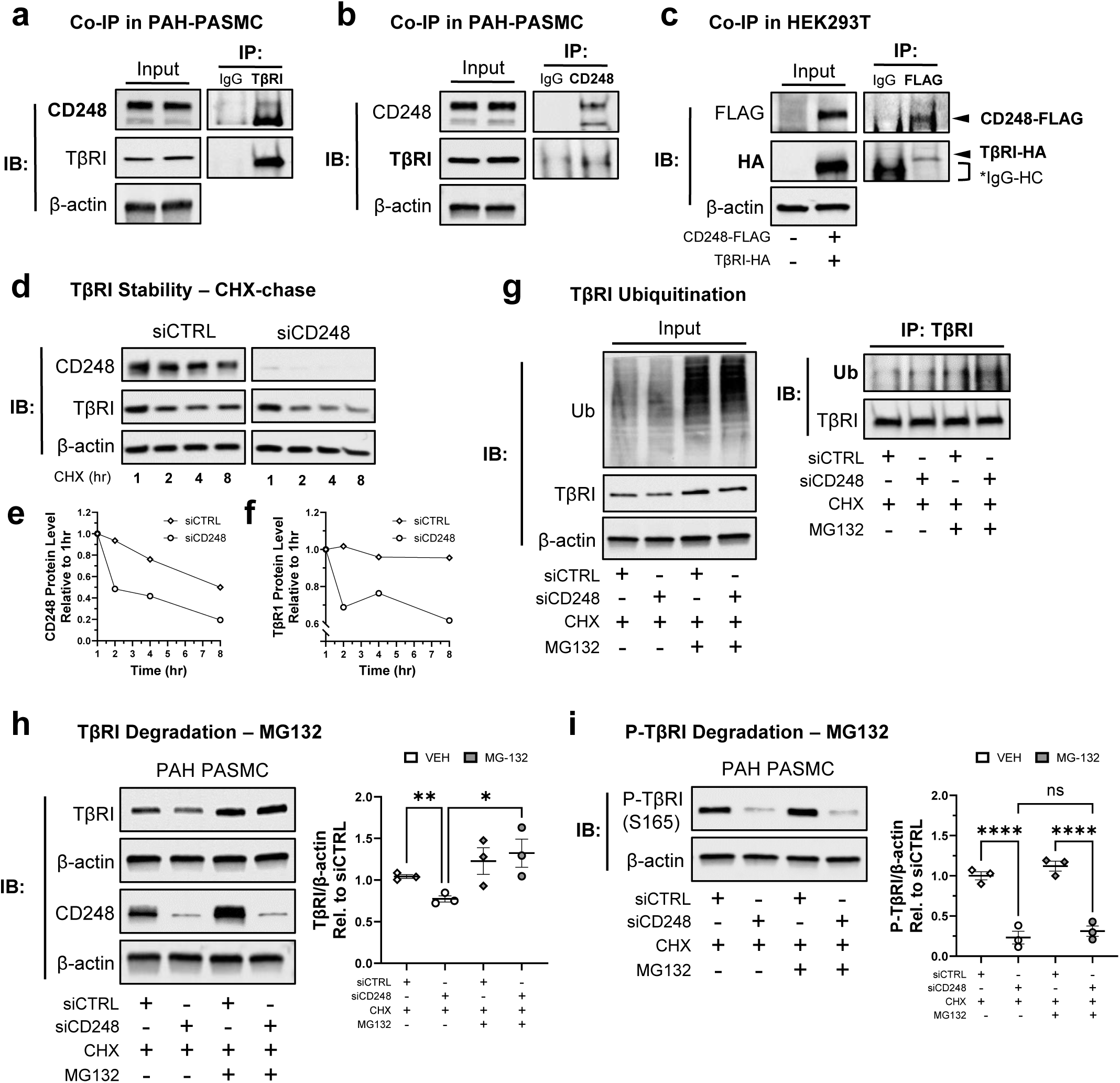
CD248 promotes TβRI stability and *de novo* phosphorylation. **(A)** Co-immunoprecipitation (Co-IP) of TβRI with CD248. TβRI was immunoprecipitated (IP) from PAH PASMC whole cell lysates followed by immunoblotting for CD248. An input lysate is shown. **(B)** Co-IP of CD248 with TβRI. CD248 was immunoprecipitated (IP) from PAH PASMC whole cell lysates followed by immunoblotting for TβRI. An input lysate is shown. **(C)** Co-IP of CD248 with TβRI in HEK293T cells. HEK293T cells were transiently co transfected with 1μg/mL of pCMV3-CD248-FLAG and pCMV3-TβRI-HA (**Supplemental Fig 7)**. Cells were collected 48 hours post-transfection followed by IP for CD248 then immunoblot for TβRI. Transfected and untransfected input lysates are shown. CD248-FLAG and TβRI-HA are indicated by arrows. IgG-heavy chain is indicated by a bracket (]). **(D)** Cycloheximide (CHX; protein translation inhibitor)-chase of TβRI following CD248 KD in PAH PASMCs. Cells were transfected according to **Fig 4B**. A time course of CHX (50μg/mL) was administered for 0-8 hours followed by harvest and degradation analysis. Representative degradation plots of CD248 **(E)** and TβRI **(F)** degradation are depicted. **(G)** Representative immunoblotting of TβRI-ubiquitination following CD248 loss. CD248 KD PAH PASMCs were delivered CHX and/or MG132 (20μM; proteasome inhibitor) for 2 hours prior to harvest. TβRI was then IP from whole cell lysates and ubiquitin was probed via immunoblot. Input lysates are shown. **(H)** Representative immunoblotting for TβRI following MG132 rescue of CD248-mediated TβRI loss. CD248-deficient (KD) PAH PASMCs were delivered CHX and/or MG132 as in **G**. CD248-mediated loss of TβRI was fully reversed by the addition of MG132. **(I)** Representative immunoblotting for P-TβRI in PAH PASMCs following MG132 addition. CD248-deficient (KD) PAH PASMCs were delivered CHX and/or MG132 as in **G**. CD248-mediated loss of P-TβRI was not modified by the addition of MG132. Data are reported as mean ± SEM; *P<0.05, **P<0.01, ****P<0.0001. Immunoblots from whole cell lysates are reported as a fractional ratio relative to β-actin as a loading control. Statistical analyses were performed via repeated measures or ordinary 2-way ANOVA with uncorrected Fisher’s LSD post-hoc comparisons tests.

### Loss of CD248 prevents *de novo* phosphorylation of TβRI, promoting its proteasomal degradation

Canonical TβRI turnover occurs via ubiquitin-mediated proteasomal degradation^48^. TβRI-ubiquitination was assessed following CD248 loss and proteasomal inhibition with MG132. TβRI ubiquitination was augmented as a consequence of CD248 KD as determined by IP (**Fig 6g**). Additionally, TβRI loss (as a function of CD248 KD) was fully rescued by administration of MG132 (**Fig 6h**). These data indicate that CD248 loss increases TβRI ubiquitination and proteasomal degradation. Interestingly, proteasome inhibition did not rescue CD248-mediated loss of P-TβRI (**Fig 6i**), suggesting that CD248 facilitates TβRI phosphorylation at S165. Taken together, these data illustrate that CD248 is essential for the stability and *de novo* phosphorylation of TβRI.

### Ontuxizumab (anti-CD248) attenuates hallmarks of vascular remodeling and promotes TβRI degradation

We evaluated the impact of a clinically relevant anti-CD248 pharmacologic intervention, Ontuxizumab, on hallmarks of vascular remodeling in patient-derived PAH PASMCs. Ontuxizumab targets the C-type lectin (CTL) domain of CD248, promoting its receptor-mediated endocytosis^49^. To confirm this mechanism, we pretreated cells with Ontuxizumab and lysosomal neutralizing agent, chloroquine (CQ), then observed Ontuxizumab-mediated CD248 intracellular aggregation (**Fig 7a**). Ontuxizumab treatment attenuated mRNA levels of pro-remodeling TGF-β-responsive genes, *Fn1, Col1a1,* and *Acta2* (**Fig 7b**). This effect was paralleled by a significant reduction in TβRI mRNA levels (**Fig 7c**), similar to our genetic KD model in **Fig 5c**. Further, we performed a CHX-chase experiment following overnight Ontuxizumab pre-treatment and identified a marked increase in the rate of TβRI degradation in Ontuxizumab-treated cells (**Fig 7e,g**). Consistent with **Fig 6d**, this effect was most prominent two hours post-CHX addition where CD248 levels were approximately 50% reduced (**Fig 7f**). Ontuxizumab-mediated TβRI loss was also rescued by MG132 treatment (**Supplemental Fig 8**), indicating that Ontuxizumab treatment enhances proteasomal degradation of TβRI. Taken together, these findings provide compelling pre-clinical evidence that pharmacologic targeting of CD248 with Ontuxizumab may reverse vascular cell remodeling in PAH PASMCs by promoting TβRI instability and proteasomal degradation.

**Figure 7.**
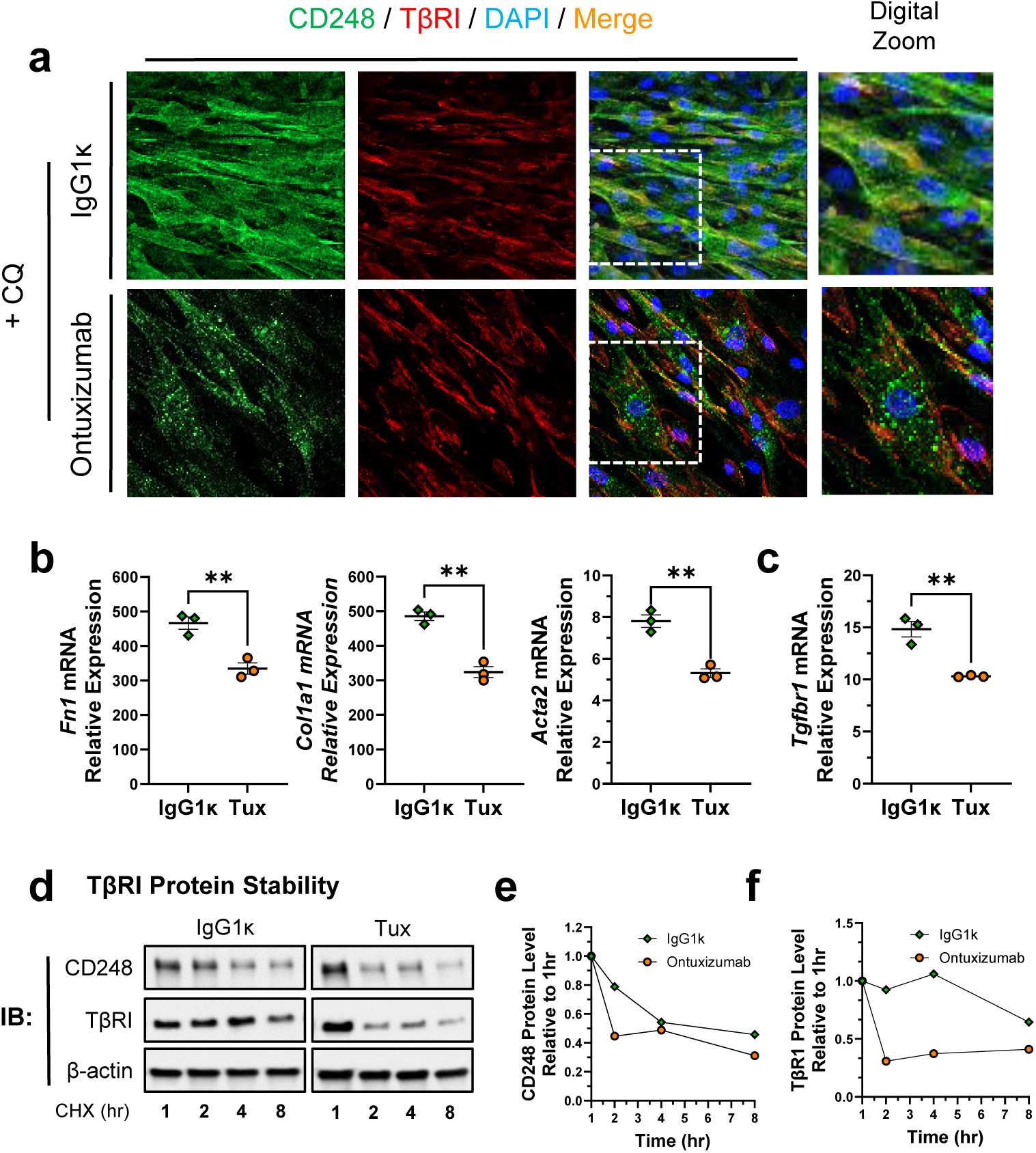
Ontuxizumab (anti-CD248) decreases vascular ECM production and TβRI stability in PAH PASMCs. (**A**) Representative IF images of CD248 (green), TβRI (red), and DAPI (blue) in PAH PASMCs treated with Ontuxizumab (anti-CD248, “Tux”; 10μg/mL) or isotype control (IgG1κ; 10μg/mL) plus chloroquine (**CQ**; **50μM**, **8hr**). Merged signal is indicated in orange. A digital zoom is indicated. (**B**) Provascular ECM remodeling target genes, *Fn1*, *Col1a1*, and *Acta2* (α-SMA) mRNA following Ontuxizumab treatment. PAH PASMCs were treated with Ontuxizumab or IgG1κ for 48 hours then mRNA expression was quantified via RTqPCR. (**C**) *Tgfbr1* mRNA following 48-hour Ontuxizumab treatment. (**D**) CHX-chase of TβRI following Ontuxizumab treatment in PAH PASMCs. Cells were treated with Ontuxizumab and rhTGF-β1 overnight. The following morning, a time course of CHX (50μg/mL) was administered for 0-8 hours followed by harvest and degradation analysis. Representative degradation plots of CD248 (**E**) and TβRI (**F**) degradation are depicted. Data are reported as mean ± SEM; *P<0.05, **P<0.01. All immunoblots are reported as a fractional ratio relative to β-actin as a loading control. Statistical analyses were performed via independent t-tests. A Mann-Whitney test was performed for nonparametric data.

## IV. DISCUSSION

TGF-β signaling is an essential regulator of vascular homeostasis^19^ that, when dysregulated, drives PAH pathogenesis^11^. The recent FDA approval of sotatercept^4^, which globally quenches ligands of this axis, marks one of the most significant advances to PAH care in decades. Emerging concerns regarding the adverse effects of global TGF-β inhibition^6, 50^ underscore the need for pulmonary vascular-specific therapies in PAH. Our work provides evidence that the transmembrane glycoprotein, CD248, is a previously unrecognized but essential co-activator of TβRI in PASMCs of the PAH pulmonary vasculature. We demonstrate in primary PAH PASMCs that CD248 knock-down attenuates cell proliferation (**Fig 4d-f**), migration (**Fig 4g**), and vascular ECM remodeling (**Fig 5f-h**). In addition, CD248 knock-out protects a preclinical murine model from H/S– induced elevation of RVSP, RV hypertrophy, and arterial muscularization (**Fig 3**). Mechanistically, CD248 interacts with and stabilizes TβRI from ubiquitin-mediated proteasomal degradation (**Fig 6d-h**). This interaction augments the phospho-activation of TβRI at S165 (**Fig 5a-b**), which ultimately enhances the downstream phosphorylation of the intracellular transduction mediator, SMAD3 at S423/425 (**Fig 5d**). Selective pharmacologic targeting of CD248 with the clinical therapeutic Ontuxizumab^38–40^ (**Fig 7a**) parallels our CD248 siRNA knock-down model, attenuating ECM deposition and promoting TβRI instability (**Fig 7d-f**). This work establishes CD248 as a novel co-activator of TβRI in PAH PASMCs. As CD248 is developmentally active but quiescent in most adult tissues, these findings suggest that anti-CD248 therapy may provide a pulmonary vascular-specific alternative to systemic TGF-β inhibition.

### CD248 expression in the pulmonary vasculature

The human CD248 gene, located on chromosome 11, uniquely lacks introns, a defining characteristic that may contribute to its transcriptional repression in adulthood and PASMC-specific expression pattern in PAH^51^. CD248 transcription is responsive to cell density, hypoxia, and glucose abundance^52–54^, all of which are essential for PAH pathogenesis^55–57^. More specifically, CD248 expression can be activated by specificity protein 1 (SP1) and hypoxia-inducible factors (HIF) 1α/2α^53^, but additional factors are certain to exist. The transcriptional reactivation of CD248 from quiescence in PAH PASMCs is beyond the scope of this manuscript but an area of active research. It is however likely that CD248 expression is tightly regulated by the complex interplay of genetic and environmental influences.

### CD248-mediated phospho-activation of TβRI

Phosphorylation of TβRI at S165 is strongly implicated in TGF-β-induced ECM deposition and cell overgrowth^58^. Interestingly, this phosphorylation event has never been described in PAH but was observed at baseline in our PAH PASMC cohort (**Fig 5a**). As pathologic TGF-β signaling is well described in PAH, constitutive phosphorylation of TβRI at S165 may provide a previously unrecognized mechanistic explanation for TGF-β hyperactivity. The TGF-β signaling cascade is first initiated by the binding of dimerized TGF-β1 ligand to TβRII, resulting in its dimerization. In turn, TβRII recruits two TβRI subunits in order to promote their phosphorylation at S165^58^. Our work is the first to report that CD248 loss confers a striking reduction in the phosphorylation of TβRI at S165 (**Fig 5a**). The role of CD248 within the activation paradigm of the TGF-β complex has never been described. We demonstrate that CD248-mediated loss of P-TβRI is likely a function of blunted *de novo* phosphorylation of TβRI rather than enhanced protein turnover or degradation (**Fig 6i**). As CD248 stabilizes TβRI (**Fig 6d-f**), it is further plausible that CD248 may increase the affinity and/or conformational interaction between TβRII and TβRI, which may augment TβRII-mediated phosphorylation of TβRI S165. In summary, we describe for the first time that CD248 is a novel regulator of TβRI phosphorylation at S165 in PAH.

### CD248-TβRI interaction

Structurally, CD248 is a single-pass transmembrane glycoprotein that includes an extracellular C-type lectin (CTL) globular domain, sushi domain, three EGF repeats, and a heavily O-glycosylated mucin-like domain^59^. Intracellularly, there is a highly conserved cytoplasmic tail with a C-terminal PDZ-binding domain that is essential for its effect on cell proliferation^60^. Here, we demonstrate that CD248 interacts with the type I TGF-β receptor, TβRI (**Fig 6a-c**). Our data suggests that the extracellular CTL domain may help mediate this interaction as Ontuxizumab, which selectively recognizes the CD248 CTL^61^, promotes TβRI instability and proteasomal degradation (**Fig 7e-h**). Ontuxizumab’s large, highly selective immunoglobulin structure may dissociate the CD248-TβRI complex, outcompeting a stable extracellular interaction between the CD248 and TβRI (presumably at the CTL domain). Our data supports this hypothesis as Ontuxizumab-treated PAH PASMCs do not show CD248-TβRI+ intracellular granules following chloroquine (lysosomal neutralization) pretreatment (**Fig 7a**). It is further plausible that CD248 interacts with TβRI in conjunction with one or more scaffolding proteins, perhaps via its C-terminal PDZ binding motif. Future work will determine the impact for both intra– and extracellular interaction domains of CD248 and TβRI. Nevertheless, these findings serve as a proof-of-principle that the pathologic CD248-TβRI complex is a pharmacologically amendable target in PAH.

### CD248 in the pulmonary vascular microenvironment

Our present study identified elevated CD248 expression via LC-MS/MS proteomic analysis of end-stage human PAH lungs where vascular remodeling hallmarks were enriched (**Fig 1d,e**). Interestingly, we observed significantly less soluble CD248 in human PAH plasma (**Fig 2f**), supporting its pulmonary vascular-specific expression pattern and attractive therapeutic potential within systemic circulation. Although it is unknown whether late-stage elevation in vascular CD248 represents sustained expression from early-to-late-stage disease, our murine studies indicate that CD248 is essential for early-stage PAH pathogenesis *in vivo* (**Fig 3**). Abundant evidence suggests that CD248 plays an essential role in vascular morphogenesis and remodeling during development and pathology^62–65^. Numerous reports also indicate that CD248 reactivation in disease is correlated with enhanced proliferation and migration^30, 31, 35, 66–68^. These processes are integrally dependent on cell interactions with ECM proteins in the extracellular microenvironment including but not limited to FN1, Col1α1, and Col4α1, all of which have been identified as ligands of the CD248 ectodomain^61, 65^. This raises the possibility that CD248 may serve as a chief mediator of matrix-to-cell mechanotransduction, which is preliminarily supported in cancer^69^.

Moreover, when CD248 is reactivated in disease, it aggravates TGF-β-mediated ECM deposition and favors a pro-remodeling microenvironment. Our data indicates that genetic loss of CD248 favors an approximately 50% or greater reduction in TGF-β-induced FN1 and Col1α1 production (**Fig 5f,h**). Furthermore, CD248 may impede the clearance of “old” matrix – largely driven by matrix metalloproteases (MMPs) – leading to an accumulation of dense, fibrotic material in the PAH lung. Our findings support this claim as CD248 loss not only impedes *de novo* ECM production but also activates the expression of the pro-remodeling matrix metalloprotease, MMP-8 (**Fig 5g**). Conversely, we report that MMP-8 levels are reduced in end-stage PAH lungs where CD248 levels are elevated (**Fig 1d**). In addition, MMP-8 KO mice have exaggerated hypoxic-induced disease^70^. As MMP-8 appears to play a vaso-protective role in PAH, anti-CD248 therapy may therefore activate these natural matrix-clearance mechanisms. Collectively, CD248 is essential for pathologic vascular remodeling in the PAH lung where its expression enhances extracellular matrisome remodeling^71^, wound healing, cell proliferation, and TGF-β-responsive pathways, all of which were elevated at baseline in our human proteomics analysis (**Fig 1e**).

In addition to its effect on ECM remodeling, we report that CD248 loss attenuates expression of α-SMA, a cytoskeletal marker of smooth muscle cell contractility (**Fig 5f,h**). Although TGF-β signaling promotes α-SMA expression, ECM composition and stiffness – via matrix-cell mechanotransduction – are also known to regulate α-SMA levels^72^. As we describe compositional reorganization of the vascular ECM following CD248 loss, it cannot be negated that α-SMA loss may be a function of both ECM– and TGF-β-mediated signals. Notably, Ontuxizumab treatment parallels these knock-down findings, reducing the expression of vascular ECM proteins, FN1, Col1α1 and α-SMA in PAH PASMCs. These findings place CD248 at the crux of vascular homeostasis in PAH. Its pathologic reactivation is essential for PASMC proliferation/migration, and its loss attenuates ECM deposition while activating the production of the vasoprotective matrix metalloprotease, MMP-8.

### Conclusion

Our study demonstrates that CD248, a transmembrane glycoprotein, is selectively activated in PASMCs of the PAH pulmonary vasculature where it enhances pulmonary vascular remodeling via co-activation of TβRI. Ontuxizumab, an anti-CD248 clinical therapeutic, exhibits compelling preclinical efficacy by attenuating hallmarks of vascular remodeling in primary human PAH PASMCs. Our findings support the potential for anti– CD248 therapy as a pulmonary vascular-specific alternative to systemic TGF-β inhibition.

## VI. ACKNOWLEDGMENTS

The authors acknowledge the UAB Pathology Core Research Lab, the Flow Cytometry and Single Cell Core Facility, the support of the Center for AIDS Research Grant AI027767, and The O’Neal Comprehensive Cancer Center Grant CA013148. We thank the UAB Center for Exercise Medicine and the UAB Medical Scientist (MD-PhD) Training Program for their financial and professional support. We thank Dr. Christian Faul, Dr. A Brent Carter, and Dr. Jennifer Larson-Casey (UAB) for their insight to this study and for providing cell and molecular resources. The LC-MS/MS proteomics analysis was performed at the Environmental Molecular Sciences Laboratory, a U.S. Department of Energy (DOE) National Scientific User Facility located at the Pacific Northwest National Laboratory (PNNL) in Richland, WA. PNNL is a multi-program national laboratory operated by the Battelle Memorial Institute for the DOE under contract DE-AC05-76RL01830.

## VIII. GRAPHICAL ABSTRACT

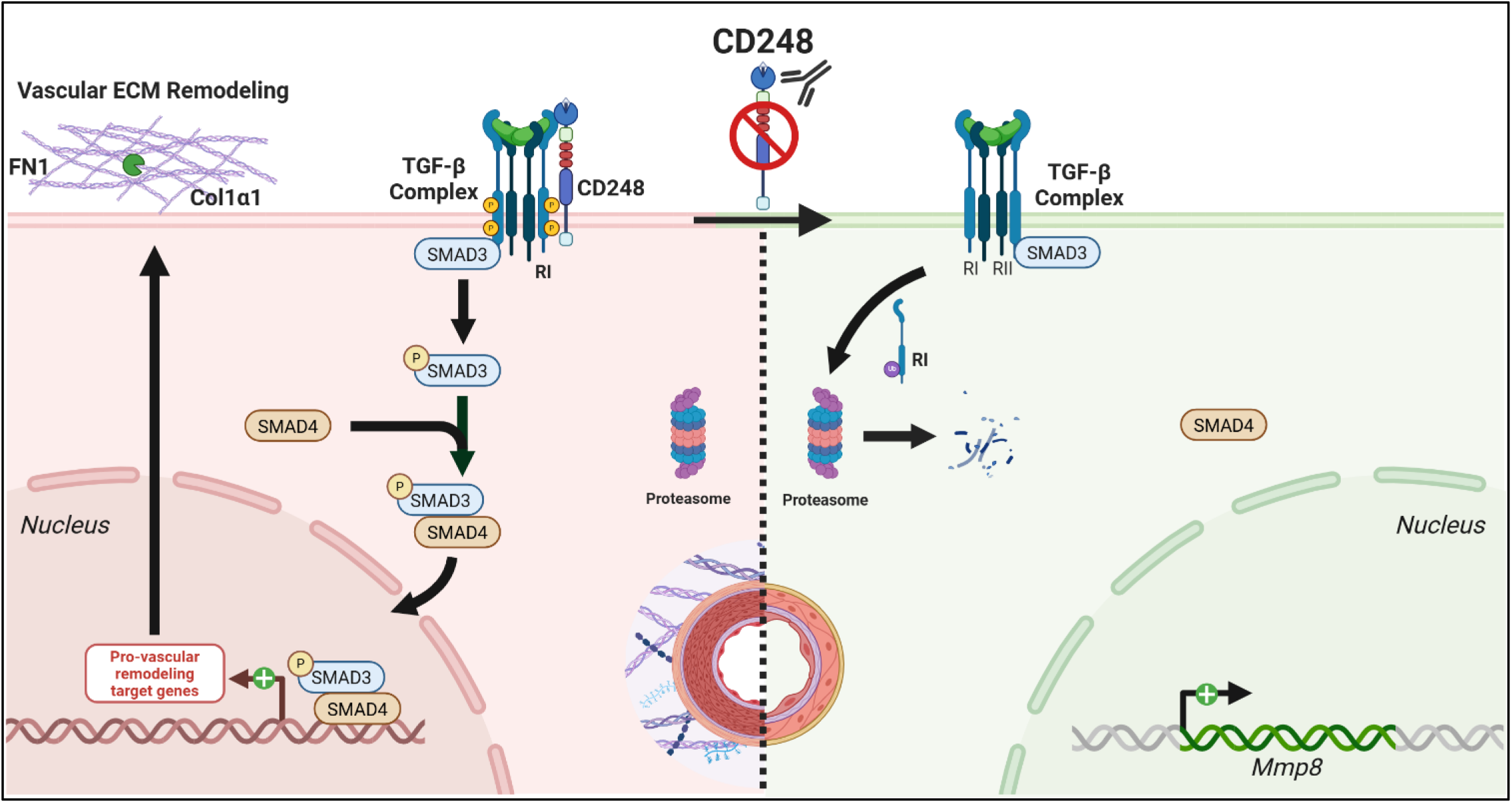

## IX. METHODS

### Human Lung Cohort and Isolation

All explanted lungs were collected either at the Cleveland Clinic through an Institutional Review Board approved protocol, or they were provided by Baylor, Stanford, Vanderbilt, University of Alabama at Birmingham, or Allegany College of Maryland under the Pulmonary Hypertension Breakthrough Initiative (PHBI). Funding for the PHBI was provided by the Cardiovascular Medical Research and Education Fund (CMREF). Human lung tissues used in this study were from 6 donor lung explants not suitable for lung transplantation and 6 idiopathic PAH patients (**Supplemental Table 1**).

### Primary Human Vascular Cells

Human PASMCs and pulmonary arterial endothelial cells (PAECs) were isolated from elastic pulmonary arteries dissected from both control and IPAH lungs obtained at explantation as previously described^73^. Briefly, human pulmonary arteries were minced and digested overnight in Hanks’ balanced salt solution containing collagenase and DNase and HEPES buffer (Sigma, St. Louis, MO). Upon removal of PAECs, smooth muscle cells (PASMCs) were released from the artery tissue, filtered with a 100-μm-pore nylon cell strainer (BD Falcon, Bedford, MA), cultured in DMEM/F-12 medium supplemented with 10% FBS (Bio-Whittaker, Walkersville, MD) and antibiotics, and incubated at 37°C, 5% CO2 with 90% humidity follow by media changes at 24 hours every 4 days until confluence. The PASMCs were confirmed routinely through positivity staining for α-smooth muscle cell actin (Sigma/Aldrich, St. Louis, MO).

### Human Plasma Collection

Blood was drawn from patients recruited from the Pulmonary Vascular Program at the Cleveland Clinic and banked in a sample biorepository. Pulmonary hypertension was confirmed by right heart catheterization. PAH categories were identified based on the updated clinical classification of pulmonary hypertension. Healthy controls were defined as patients with no history of pulmonary/cardiac disease or symptoms. All participants signed a consent form that was approved by the Cleveland Clinic Institutional Review Board before participation in the study.

### LC-MS/MS Proteomics

Protein Extraction and Sample Preparation for Proteomics Analysis. All chemicals for the proteomics sample preparation were obtained from Sigma-Aldrich (St. Louis, MO) with the following exceptions: dithiothreitol, bicinchoninic acid protein assay (BCA), iodoacetamide, and TMTpro 16plex label reagent set were obtained from Thermo Fisher Scientific, Trypsin was obtained from USB Thermo Fisher Scientific. Ultrapurified water was obtained from a Millipore Milli-Q system. Extraction of the proteins was performed using the MPLEx method on pieces of lung tissue^74^. 940 µL of ice-cold 3:4 methanol:water was added to each sample in a 5.0 mL Eppendorf tube. The sample was then homogenized using an Omni tissue homogenizer (Omni, Kennesaw, GA) with a disposable tip. Following homogenization, 1,060 µL of ice-cold chloroform was added to each sample, and the tubes were vortexed for 1 min. All samples were placed on ice for 5 min followed by another 1-min vortex. The samples were then centrifuged at 10,000 x *g* for 10 min at 4°C. An interlayer protein disc (located between the polar and apolar layer) was harvested, rinsed with methanol, and resuspended in 50 mM ammonium bicarbonate using vortexing and a brief sonication. A Pierce BCA protein assay was performed to determine the approximate protein concentration of each sample. 500 µg of protein were utilized from each sample, with volumes normalized using denaturing buffer (50 mM ammonium bicarbonate containing 8M Urea). After 5 min at 95°C, Dithiothreitol was added to a 5 mM concentration, and the samples were incubated on a Thermomixer (Eppendorf) for 1 h at 37°C with shaking at 850 rpm. Iodoacetamide was added to a 40 mM concentration, and the samples were incubated as before but in darkness. Samples were diluted eightfold with 1.14 mM CaCl_2_ in 50 mM ammonium bicarbonate, and trypsin was added in a 1:50 trypsin:protein ratio. The samples were incubated for 3 h at 37°C (vertical inversion/rotation) at about 20 rpm. Resulting peptides were desalted using a solid phase extraction (SPE) using 1 mL/50 mg C18 columns from Phenomenex following supplier instructions. Briefly, columns were conditioned with 2 mL of methanol followed by 3 mL of 0.1% trifluoroacetic acid (TFA) in water. A sample was applied to each column followed by 4 mL of 95:4.9:0.1 H_2_O:MeCN:TFA. Samples were eluted into 1.5 mL microcentrifuge tubes with 80:19.9:0.1 MeCN:H2O:TFA and concentrated in a vacuum concentrator to 50 µL. Another BCA protein assay was performed to obtain peptide concentration. For tandem mass tag (TMT) labeling, 100 µg of peptides from each sample were aliquoted, dried, and reconstituted to a 5 µg/µL concentration with 50 mM HEPES, pH 8.5. TMT reagent (20 mg/mL) was added to the peptides in a 1:1 ratio (wt:wt). Samples were incubated at 25°C for 1 h at 400 rpm in a Thermomixer. Samples were then diluted to 2.5 µg/µL with 50 mM HEPES pH 8.5, 20% acetonitrile. The samples labeled with different TMT reagents were combined into one tube, diluted with water to reduce the acetonitrile concentration to less than 5%, and desalted using C18 solid phase extraction (SPE). The multiplexed sample was concentrated to 100 µL, and a BCA protein assay was again performed for quantitation. The peptide mixture was then fractionated using high pH reverse phase chromatography on a Waters XBridge column and concatenated into 12 fractions as previously outlined^75^. Each fraction was concentrated on a vacuum concentrator to 50 µL, the peptides were then quantitated and diluted with water to 0.07 µg/µL for LC-MS analysis.

LC-MS/MS Proteomics. The LC was configured to load the sample first on an SPE column followed by separation on an analytical column. Analytical columns were made in-house by slurry packing 3 μm Jupiter C18 stationary phase (Phenomenex) into a 70 cm long, 360 μm OD × 75 μm ID fused silica capillary tubing (Polymicro Technologies). Samples were loaded on the SPE column via a 5 μL sample loop for 30 min at a flow rate of 5 μL/min and then separated by the analytical column using a 120 min gradient from 99% mobile phase A (MP-A) to 5% MP-A at a flow rate of 300 nL/min. MS analysis was started 20 min after the sample was moved to the analytical column. After the gradient was completed, the column was washed with 100% mobile phase B (MP-B) then reconditioned with 99% MP-A for 30 min. The effluents from the LC column were ionized by electrospray ionization by applying 2,200 V to the metal union between the column and the electrospray tip. Electrosprayed ions were introduced into the mass spectrometer (Orbitrap Q Exactive HFX) via a heated capillary (5.8 cm long with a rectangular slit of 1.6 mm long and 0.6 mm wide) maintained at 250°C for ion desolvation. The resulting ions were mass analyzed by the Orbitrap at a resolution of 60,000 covering the mass range from 300 to 1,800 Da with a maximum injection time of 20 ms and automated gain control (AGC) setting of 1E5 ions. Mass spectra were recorded in profile mode. The most abundant ions were subjected to MS2 analysis using the top speed mode, acquiring the top 12 ions in the cycle time. The parameters used for these analyses were as follows: For MS2, ions were isolated by quadrupole mass filter in monoisotopic peak selection mode using isolation window of 0.7 Da, maximum injection time of 100 ms with AGC setting at 5E3 ions, and fragmented by high-energy collision dissociation (HCD) with nitrogen at 30% normalized collision energy. Fragment ions were mass analyzed by the Orbitrap at a resolution of 30,000, and spectra were recorded in the centroid mode. Ions once selected for MS2 were dynamically excluded for the next 45 s. Mass spectra were recorded in centroid mode. The instrument raw files are publicly available on MassIVE (Server: massive.ucsd.edu).

Proteomics Data Analysis. Raw MS data were converted to peak lists (DTA files) using the DeconMSn^76^ and searched with MS-GF+^77^ against the human UniProt Reference Proteome (UP000005640, downloaded in July 2021), and general contaminants including bovine trypsin. The identified spectra were filtered based on their MS-GF+ scores (5.4693e-11), and only the proteins with two proteospecific peptides were conserved, resulting in a PSM, peptide, and protein false discovery rate (FDR) <1%. Peptide and protein rollup were performed by summing spectra intensities for the reporters. Statistical analyses were performed using in-house RomicsProcessor v1.1.0 (R package, https://github.com/PNNL-Comp-Mass-Spec/RomicsProcessor, release ID 72d35c9). Briefly, the data were imported as a multilayered R object with its associated metadata. The intensities were then log2 transformed and filtered to allow maximal missingness of 70% within at least one given group. After median normalization, the missing values were imputed using a previously described method using a random downshifted distribution^78^. A principal component analysis (PCA)-based dimension reduction analysis was performed. ANOVA and two-tailed heteroscedastic Student’s t-test were used to compare the experimental groups (CTRL and PAH). Pathway enrichment analyses of differentially expressed proteins were performed using Metascape^41^ (v3.5.20260201)

### Hypoxia-Sugen (H/S) Mouse Model

10-12-week C57BL/6 mice were purchased from Jackson Laboratories (Bar Harbor, Maine, USA). Male mice were used in this study as females fail to develop significant pathology in the hypoxia-sugen model, an established model of murine pulmonary hypertension^44^. CD248 null mice (CD248^−/−^) have been previously described^79^ and were provided for this study by the National Cancer Institute (St. Croix Laboratory) via an approved MTA and were expanded and maintained within the UAB Hugh Kaul Animal Resource Program Facility. Genotypic confirmation of CD248 knock-out was performed using the following primer sequences:

CD248^−/−^ Forward – GCCCCAGCCCACTCCTTAC

CD248^−/−^ Reverse – GCAACCCAATCCATAGCAGC

CD248^WT^ Forward – CTTGGGTGGAGAGGCTATTC

CD248^WT^ Reverse – AGGTGAGATGACAGGAGATC

Animals were housed under pathogen-free conditions with food and water *ad libitum*. Littermates were randomly assigned to experimental groups. *Model generation:* Mice assigned to the H/S arm of the study received hypoxia (10% O_2_) (Chamber: [model 110, BioSpherix, Redfield, New York) for 3 weeks with or without weekly subcutaneous (SQ) injection of SU5416 (20mg/kg in DMSO). Alternatively, control mice were maintained at room air (21% O_2_) and received weekly subcutaneous vehicle injections. At 3 weeks of H/S or control conditions, all mice were assessed for hemodynamic measurements and cardiac pathology. Following these endpoint procedures, lung and cardiac tissues were collected for molecular analyses. *Vascular wall thickness:* An average of 10 H&E vascular lung fields (average vessel diameter <150 μm) was used to calculate the diameter (d) of the circle created by the outer (O) and inner vascular wall (I) that produced area (A) according to the following: 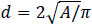. Wall area relative to total vessel area was calculated as a normalized ratio of diameters 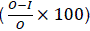. All experiments and procedures were approved by the University of Alabama at Birmingham Institutional Animal Care and Use Committee (IACUC).

### Hypoxia-Sugen Rat Model

All animal experiments were performed under the approval of the University of Alabama Institutional Animal Care and Use Committee (IACUC) and in accordance with the National Institutes of Health Guide for the care and use of laboratory animals. This manuscript adheres to the ARRIVE guidelines. In brief, PAH was established in adult male Sprague Dawley rats (160-200g) as previously described^44, 80^. Rats were injected subcutaneously with the VEGFR2 inhibitor SU-5416 (20 mg/kg; Sigma) and exposed to normobaric hypoxia (10% O2) for 3 weeks in a hypoxic chamber (BioSpherix), equipped with an oxygen controller (ProOx 360). At the end of 3 weeks, RVSP was measured to confirm model establishment, as previously described^80^. Control animals were maintained in room air (21% O2). The number of animals used in each experiment is indicated in the figure legends. All experiments were carried out independently for a minimum of two times.

### Hemodynamic Measurements

Intact right heart hemodynamic analysis was performed in mice using a pressure-volume catheter. Animals were anesthetized using 2% isoflurane gas. Depth of anesthesia was monitored by observing chest wall movement, heart rate, blood pressure, muscle tone, and stimulus perception. The right internal jugular vein (IJV) was first dissected then cannulated with a 30-gauge needle. A 4-electrode Millar Mikro-Tip pressure-volume catheter (SPR-1000; AD Instruments) was then inserted into the right IJV and advanced into the right ventricle (RV) where stable systolic pressure (RVSP) tracings were recorded. The pressure catheter was then removed and immediate ligation of the IJV proximal and distal to the cannulation site preceded closure of the incision with surgical adhesive.

### Fulton Index (FI)

Immediately following 2-point euthanasia, we excised the heart from each animal. While submerged in PBS, we removed all remnants of the great vessels, surrounding fascia, and right/left atria, leaving the right (RV) and left (LV) ventricles intact. The RV was then carefully dissected from the LV using precision snips. Following thorough drying, the RV and the LV + septum (LV+sep) were weighed with a precision scale (Ohaus AX224). Fulton indices (RV/LV+sep) were calculated for all mice hearts to assess right ventricular hypertrophy.

### Lung Homogenization

Lungs were separated by lobes into RINO tubes (Cat #NAVYR1-RNA; Next Advance) and filled with ice cold PBS + 2X HALT protease and phosphatase inhibitors (ThermoScientific 78442) and DNase I (IBI XH28701). Flash-frozen lung sections were added then homogenized three times using a bullet blender (Bullet Blender 24; Next Advance) at max speed for 1 minute at 4°C. The cell suspension was then aliquoted and frozen at –80°C.

### General Eukaryotic Cell Culture

Primary CTRL and PAH pulmonary artery smooth muscle (PASMC) and endothelial cells (PAECs) were obtained from the pulmonary hypertension breakthrough initiative (PHBI) and maintained in liquid nitrogen. Passage 4-9 cells were seeded on 10 cm dishes at an average density of 1×10^6^ cells in vascular SMC media (Lifeline Technologies; LS-1040). Experiments were performed in 6-well plates (cat no: 353046, Corning) at an average seeding density of 2×10^5^ cells. All plates were harvested at 80-90% confluence prior to downstream analyses.

### PASMC siRNA Knock-Down (KD)

150-200k PASMCs were plated per well in 6-well plates. Following adhesion, cells were transfected in Opti-MEM™ (Gibco #11058021) according to manufacturer’s instructions using Lipofectamine™ LTX (cat no: 15338030; ThermoFisher) + 40nM siRNA (Dharmacon; constructs below). Transfection media was changed to vascular SMC media following overnight (16-18 hours) transfection. Cells were grown for an additional 48 hours prior to protein/RNA expression analyses.

siRNA Constructs:

a. siCD248 (SMARTpool) Target Sequences:

1. GCGCAUCACUGACUGCUAU
2. GCAGCCAACUAUCCAGAUC
3. GGACCUCGGAGAUGAGUUG
4. CCACCAGCCUCCUGUGAUC
b. siCTRL Target Sequences:

Sense: UAAGGCUAUGAAGAGAUACUU

Anti-Sense: GUAUCUCUUCAUAGCCUUAUU

### Click-iT® EdU Incorporation

Cells were transfected with CD248 targeted or untargeted siRNA overnight then grown for 48 hours to 80-90% confluence. 24 hours prior to EdU incorporation, cells were supplemented with 20% FBS. Click-iT® EdU Incorporation was performed the following morning according to manufacturer’s instructions (Invitrogen; C10632). In brief, transfected cells were treated with 10μM EdU for 4 hours at 37°C. Cells were then trypsinized and washed with 1% BSA in PBS. Cells were fixed (4% PFA in PBS) then permeabilized for 15 minutes at each step. The Click-iT® reaction cocktail (AF488-congugated picolylazide) was added for 30 minutes at room temperature then cells were washed prior to flow cytometry analysis. A BD FACSymphony A5 Analyzer was used to detect EdU-AF488+ single cells. The gating strategy for all experiments is included in the supplemental material.

### MTT Assay

Cells were transfected with CD248 targeted or untargeted siRNA overnight then grown for 48 hours to 80-90% confluence in 6-well plates. Cells were trypsinized and counted. Equal cell numbers between groups were then aliquoted and treated with MTT reagent (1:10) for 2 hours at 37°C. Following brief centrifugation, the MTT reagent was solubilized with DMSO. Absorbance (570nm) of each sample was measured using a BioTek SynergyMx plate reader.

### Wound Healing Assay

Cells were transfected with CD248 targeted or untargeted siRNA overnight then grown for 48 hours to >90% confluence. A P200 pipette tip was used to create a horizontal scratch wound across each well. Cell debris was aspirated, cells were washed 1X with PBS, and fresh SMC media was carefully applied prior to imaging. Images of each wound were captured at baseline and at 24 hours using a Nikon Eclipse Ts2 microscope. Serial images were quantified for % wound closure using an optimized ImageJ plugin^81^.

### TGF-β1 Stimulation in PASMCs

Cells were transfected with CD248 targeted or untargeted siRNA overnight then grown for 48 hours to 80-90% confluence. For acute TGF-β1 stimulation, transfected cells were treated with either recombinant human TGF-β1 (rhTGF-β1; 10ng/mL; Peprotech #100-21) or DMSO for 30 minutes. For prolonged TGF-β1 stimulation, transfected cells were grown for 24 hours to 60-70% confluence. rhTGF-β1 (10ng/mL) or DMSO (vehicle) was delivered for 48 hours prior to downstream analyses.

### Cycloheximide Protein Translation Inhibition

Cells were transfected with CD248 targeted or untargeted siRNA overnight then grown for 48 hours to 80-90% confluence. At 48 hours post-transfection, cells were treated with cycloheximide (CHX; 50μg/mL; Millipore) and harvested at 0-, 0.5-, 1-, 2-, 4-, and 8-hour time points. For pharmacological inhibition, cells were grown to 70-80% confluence then treated with Ontuxizumab (10μg/mL; MCE #HY-P99778) or isotype control (cat no; MCE #HY-P99001) and rhTGF-β1 (10ng/mL) overnight. The following morning, cells were treated with CHX (50μg/mL; 0-8 hours) and harvested for endpoint assessments.

### MG132 Proteasome Inhibition

Cells were transfected with CD248 targeted or untargeted siRNA overnight then grown for 48 hours to 80-90% confluence. At 48 hours post-transfection, cells were treated with CHX (50μg/mL) alone or CHX + MG132 (20μM) for 2 hours prior to collection and downstream analyses. For pharmacologic inhibition, cells were grown to 70-80% confluence then treated with Ontuxizumab (10μg/mL) or isotype control (10μg/mL) and rhTGF-β1 (10ng/mL) overnight. The following morning, cells were treated with CHX (50μg/mL) alone or CHX + MG132 (20μM; Millipore #474791) for 4 hours prior to collection and downstream analyses.

### Bacterial Transformation

*E. coli* strain DH5α (Cat #C2987H; New England Biolabs) cells were utilized for plasmid transformation and amplification. DH5α cells were transformed with 10ng of each respective plasmid then incubated on ice for 20 minutes. Each sample was then heat shocked at 42°C for 45 seconds followed by 2 minutes of recovery on ice. Fresh Luria Broth (LB) (Cat #IB49020; IBI Scientific) was then added to each sample followed by 45 minutes of orbital shaking at 37°C. Transformed cultures were plated for selection on LB agar + kanamycin. Single colonies were selected the following morning for overnight (ON) expansion in LB broth + kanamycin. Selected ON cultures were frozen at –80°C in 25% glycerol. Minipreps of ON selection cultures were performed according to manufacturer’s instructions (Cat #T1110S; New England Biolabs) to purify plasmid.

### HEK293T Transient Overexpression

HEK293T cells were graciously provided by Dr. Christian Faul (UAB; Birmingham, AL). HEK293T cells were plated in 6-well dishes to approximately 50% confluence. Cells were transfected according to manufacturer’s instructions using Lipofectamine™ LTX (Cat #15338030; ThermoFisher) plus varying concentrations of plasmid (0-2μg). Following ON transfection, HEK293T cells were changed to fresh media then grown for an additional 24-48 hours prior to harvest for downstream analyses.

### Western Blotting

Adherent cells were collected by scraping and centrifugation then lysed in RIPA buffer (Cat #9806; Cell Signaling) + HALT protease and phosphatase inhibitors. Protein concentration was measured with a Bradford protein assay following centrifugation. All samples were prepared in reducing conditions using Laemmli buffer (#1610747; Bio-Rad) and β-mercaptoethanol. Samples were denatured at 95°C prior to gel electrophoresis. 20-30μg of protein per sample were separated by SDS-PAGE using 4-20% precast Ready Gels (Bio-Rad) then transferred to nitrocellulose (0.45um; Amersham #10600003) or PVDF (0.22um; Amersham #10600021) membranes. Membranes were washed with Tris-buffered saline + 0.1% Tween 20 (cat no: BP337-500; Fisher Scientific) (TBST) then blocked with 5% nonfat milk or BSA. Primary antibodies were incubated overnight at 4°C or for 2 hours at room temperature. Primary antibodies used in this study are listed in the table below. Following primary antibody incubation, membranes were washed 3X in TBST. Membranes were then incubated with species– and isotype-specific, HRP-conjugated secondary antibodies for 1 hour at room temperature. Following 3 final washes in TBST, enhanced chemiluminescence detection solution was used to develop all membranes. A GE Imaging System (GE Healthcare, USA) was used to capture all images and densitometric analyses were performed using ImageJ-FIJI software.

Primary Antibodies:

| Antibody | Species | Isotype | Company | Cat # | Block | Dilution |
| --- | --- | --- | --- | --- | --- | --- |
| **CD248** | Ms | IgG | Proteintech | 60170-1-Ig | 5M | 1:1000 |
| **PLXDC1** | Ms | IgG | Novus Bio | NB100-56557 | 5M | 1:1000 |
| **POSTN** | Rb | IgG | Novus Bio | NBP1-30042 | 5M | 1:1000 |
| **Col1α1** | Rb | IgG | Abcam | ab138492 | 5M | 1:1000 |
| **Col1α1** | Rb | IgG | Abcam | ab260043 | 5M | 1:1000 |
| **Col3α1** | Rb | IgG | Abclonal | A3795 | 5M | 1:1000 |
| **FN1** | Ms | IgG | Santa Cruz | sc-8422 | 5M | 1:1000 |
| **α-SMA** | Ms | IgG | Invitrogen | MA5-11547 | 5BSA | 1:1000 |
| **α-SMA** | Rb | IgG | Invitrogen | PA5-85070 | 5BSA` | 1:1000 |
| **PCNA** | Ms | IgG | Invitrogen | MA5-11358 | 5M | 1:1000 |
| **TGFBRI** | Rb | IgG | Invitrogen | PA5-98192 | 5M | 1:5000 |
| **TGFBRII** | Ms | IgG | Santa Cruz | sc-17792 | 5M | 1:500 |
| **β-actin-HRP** | Ms | IgG | Millipore | A1978 | 5M | 1:5000 |
| **P-SMAD3** | Rb | IgG | Cell Signaling | 9520S | 5M | 1:800 |
| **SMAD3** | Rb | IgG | Cell Signaling | 9523S | 5M | 1:900 |
| **FLAG (DDK)** | Rb | IgG | Cell Signaling | 3724S | 5M | 1:1000 |
| **HA** | Rb | IgG | Cell Signaling | 3724S | 5M | 1:1000 |
| **P-TGFBRI** | Rb | IgG | Invitrogen | PA5-40298 | 5M | 1:2000 |
| **Ubiquitin** | Ms | IgG | Cell Signaling | 3936S | 5M | 1:1000 |

HRP-conjugated Secondary Antibodies:

– GtαRb IgG-HRP (cat no: 31466; ThermoFisher)
– GtαMs IgG-HRP (cat no: 31430; ThermoFisher).

### Immunofluorescence (IF)

a) Lung Tissue: Paraffin embedded lung tissues were deparaffinized by incubating in a 95°C oven for 2 hours then immediately washed in Clear-Rite 3™. Following 3 washes in Clear-Rite 3™ then slow rehydration with dilute EtOH, antigens were retrieved with 1X citrate buffer (Cat #H-3300; Vector Labs). Slides were washed 3 times with PBS then permeabilized with 0.1% Triton X-100 in PBS. Permeabilized sections were blocked with 5% BSA then primary antibodies were incubated ON at 4°C or at room temperature (RT) for 2 hours. Primary antibodies with dilutions used for IF are listed below. Following primary antibody staining, slides were washed 7X with PBS then fluorescent secondary antibodies, listed below, were incubated at room temperature for 45 minutes. Seven additional PBS washes were used to wash away excess secondary antibody. Prior to mounting, auto-fluorescence quenching solution (#SP-8400-15; Vector Labs) was applied to each slide followed by NucBlue nuclear stain. IF images were captured using a Nikon fluorescent microscope.
b) PASMC: Cells were cultured to 80-90% confluence, fixed with 4% PFA, washed 3X with PBS, then permeabilized with PBS + 0.1% TritonX. Slides were blocked with BSA and stained with primary antibodies for 2 hours at room temperature (RT). Following 3 PBS washes, secondary antibodies were added for 1 hour at RT. All slides were washed 3-4X with PBS, stained with NucBlue, then mounted for confocal microscopy (Zeiss).

Primary Antibodies:

| Antibody | Species | Isotype | Company | Cat # | Block | Dilution |
| --- | --- | --- | --- | --- | --- | --- |
| **CD248** | Ms | IgG | Proteintech | 60170-1-Ig | 5BSA | 1:250-500 |
| **TβRI** | Rb | IgG | Invitrogen | PA5-98192 | 5BSA | 1:250 |
| **α-SMA-Cy3** | - | - | Sigma | C6198 | 5BSA | 1:250 |

Fluorescent Secondary Antibodies:

| Host | Species | Isotype | Fluorophore | Company | Cat # | Dilution |
| --- | --- | --- | --- | --- | --- | --- |
| Chicken | Ms | IgG | AF488 | Invitrogen | A21200 | 1:2000 |
| Donkey | Rb | IgG | AF594 | Invitrogen | A21207 | 1:2000 |
| Goat | Ms | IgG | AF700 | Invitrogen | A21036 | 1:2000 |

### Immunoprecipitation (IP) and co-IP

Cells were collected by centrifugation then lysed in RIPA or non-ionic lysis buffer (1% NP-40 in PBS) + HALT protease and phosphatase inhibitors. A 1-10% lysate was saved for each sample as input material and prepared as above (see *Western Blotting*). Remaining lysates were precleared with protein A/G beads (sc2003; Santa Cruz), tumbling end-over– end for 1 hour at 4°C. Precleared lysates were incubated with primary antibodies (1-2μg) tumbling end-over-end at 4°C ON. The following day, A/G capture beads were added with end-over-end tumbling for 2 hours at 4°C. Capture beads were then washed 2X with ice-cold non-ionic lysis buffer, 4X with detergent buffer (75mM Tris, 150mM NaCl, 1% Triton X-100), and 3X in 75mM Tris buffer. Protein-bead complexes were then dissociated at 95°C in 4X Laemmli + β-mercaptoethanol.

### Gene Expression Analyses

Total RNA extraction (ThermoScientific #K0732; IBI Scientific #IB47302;) from PASMCs and mouse lung homogenates, cDNA synthesis (Cat #K1652; ThermoFisher), RT-qPCR, and gene expression analyses were performed as previously described^82, 83^. In brief, RNA extraction from PASMCs or mouse lung homogenates was performed via RNA precipitation in EtOH and column-based purification according to manufacturer’s instructions. Quantity and quality of RNA was assessed via Nanodrop (ThermoFisher). cDNA was synthesized according to manufacturer’s instructions. For gene expression analysis, 10ng of cDNA was prepared for real time quantitative PCR (RT-qPCR) (StepOnePlus Real-Time PCR; ThermoFisher) using the below TaqMan™ probes. (Life Technologies/Applied Biosystems). Relative mRNA expression was calculated using the 2^-ΔCT^ method.

TaqMan™ Probes: *Cd248* (Hs00535586), *Fn1* (Hs01549976), *Col1a1* (Hs00164004), *Acta2* (Hs00426835), *Mmp8* (Hs01029057).

### Quantification and Statistical Analysis

Statistical analyses for each experiment can be found in their respective figure legends. Western blot images were quantified using ImageJ-FIJI and densitometric comparisons are reported relative to β-actin loading control. All RT-qPCR experiments were performed in no less than biological triplicate. Technical duplicates/triplicates of all biological replicates were analyzed for relative expression by calculating the 2^-ΔCT^ relative to GAPDH expression. Statistical analyses were performed using GraphPad Prism 10. In brief, independent Student’s t-tests were used to analyze normally distributed (Shapiro-Wilks p>0.05) 2-group comparisons. A Mann-Whitney test was used to analyze 2-group comparisons that were not normally distributed. Paired t-tests were used to compare 2-group treatments effects across multiple human subjects. For normally distributed data with >2 groups and 1 independent variable, a one-way ANOVA was performed with Tukey’s post-hoc comparisons; non-normal data was analyzed with a Kruskal-Wallis test. Data containing >2 groups and 2 independent variables was analyzed with a 2-way ANOVA with Tukey’s post-hoc comparisons. Data are reported as mean ± standard error of the mean (SEM). Outliers were determined via ROUT or Grubbs’ tests. Statistical significance was represented as follows: *p<0.05, **p<0.01, ***p<0.001, ****p<0.0001.

